# Nanoscale organization of betaII-spectrin within segments of the membrane-associated periodic skeleton in mouse sciatic nerve axons

**DOI:** 10.64898/2026.02.19.705149

**Authors:** Nahir Guadalupe Gazal, Gonzalo Escalante, Lucía F. Lopez, Laurence Goutebroze, Mariano Bisbal, E. Axel Gorostiza, Alan M. Szalai, Fernando D. Stefani, Nicolás Unsain

**Author notes:** These authors contributed equally to this work (Nahir Guadalupe Gazal, Gonzalo Escalante).

## Abstract

The actin/spectrin membrane-associated periodic skeleton (MPS) is a ubiquitous cytoskeletal structure essential for axonal integrity and function. Primarily studied in cultured neurons, the MPS has been extensively modeled as actin rings spaced by spectrin tetramers, the latter assumed to be regularly and densely distributed across the axonal perimeter. However, its nanoscale organization within native tissue environments remains poorly understood. In this study, we investigated the three-dimensional organization of βII-spectrin in the mouse sciatic nerve using 3D-dSTORM and STED super-resolution microscopy on thin transversal cryosections. By implementing a custom quantitative analysis pipeline, we resolved the sub-diffraction architecture of the MPS across myelinated axons of diverse diameters. We confirm that βII-spectrin is localized to the inner face of the axonal plasma membrane and maintains a longitudinal periodicity of approximately 170 nm, consistent with previous observations. Crucially, 3D-dSTORM revealed that βII-spectrin along the axonal perimeter is organized in discrete nanoscale clusters with a median effective radius of 25 nm, compatible with the size of an individual spectrin tetramer visualized by indirect immunolabeling. The number of these clusters scales linearly with the axonal perimeter, maintaining a constant membrane occupancy of ∼20% across varying axon diameters. Moreover, these clusters exhibit a non-random spatial distribution with a characteristic center-to-center nearest-neighbor distance of ∼200 nm. These findings challenge simplified models of the MPS based on cultured systems and demonstrate that the MPS in peripheral nerves is composed of discrete structural units. This modular, dispersed organization may provide the structural flexibility required to withstand the mechanical demands of the peripheral nervous system while maintaining a stable periodic scaffold.

## Introduction

The actin/spectrin membrane-associated periodic skeleton (MPS) is a highly organized cytoskeletal structure found in neurons, primarily composed of actin, spectrin and adducin. This structure forms a submembrane periodic lattice along the axonal shaft, with F-actin rings separated by ∼185 nm-long spectrin tetramers. Remarkably, the MPS is ubiquitous across a wide range of neuronal types and animal species (D’Este et al., 2016; He et al., 2016). This ubiquity has motivated a thorough revision of neurobiological mechanisms to evaluate the functional implications of this structure. It has been shown that the MPS plays critical roles in the organization of membrane components (Rentsch et al., 2024), the regulation of axonal diameter (Leite & Sousa, 2016; Costa et al., 2020), the resistance to pressure and tension (Fan et al., 2017; Dubey et al., 2018), the control of microtubule dynamics (Qu et al., 2017), the organization of signaling complexes at the plasma membrane (Zhou et al., 2019), and axon fragmentation during developmental pruning (Unsain, Bordenave, et al., 2018; Wang et al., 2019).

Because its characteristic periodic features exist below the diffraction limit of light, the discovery and subsequent investigation of the MPS have relied on super-resolution microscopy techniques, including dSTORM (stochastic optical reconstruction microscopy) (Xu et al., 2013) and STED (stimulated emission depletion) (D’Este et al., 2015). Later, expansion microscopy (Martínez et al., 2020) and structured illumination microscopy (Qu et al., 2017) have also been applied. Probably due to the added technical challenges of using super-resolution techniques in tissue sections, the vast majority of studies have been conducted in neurons grown in culture. Consequently, most current knowledge arises from relatively immature axons grown in artificial 2D environments, and a significant gap exists in our understanding of how this skeleton is organized in nerve tissue, despite its relevance in neurobiology. Some studies have of course confirmed the periodic distribution of actin and spectrin in axons in tissue including mouse brain sections (Xu et al., 2013), *Drosophila melanogaster* brain (He et al., 2016) and longitudinal sections of the sciatic nerve by STED (D’Este et al, 2016). However, because cell-cell and cell-matrix interactions strongly influence cell shape and membrane components, it is likely that the structure of the MPS adopts a unique organization in intact tissue.

Spectrins can associate with the plasma membrane either directly (Grzybek et al., 2006) or indirectly through their interaction with membrane ankyrins (Bennett & Stenbuck, 1979). They also bind to actin filaments, both directly and via intermediary proteins (Cohen et al., 1980; Mische et al., 1987; Gimm et al., 2002). These interactions are further regulated by post-translational modifications affecting spectrin and its associated partners (Machnicka et al., 2014; Bennett & Lorenzo, 2016). Given its functional complexity, the spectrin tetramer is expected to play a central role in MPS function, and it is therefore unsurprising that spectrin mutations in humans lead to a range of neurological disorders (Liu & Rasband, 2019; Lorenzo et al., 2023). Current working models of the structure of the MPS typically propose that spectrin tetramers that connect adjacent actin rings exist isolated from one another, and are distributed equidistantly along the axonal perimeter - see for example schemes in (Xu et al., 2013; Leite et al., 2016; Fan et al., 2017; Zhang et al., 2017; Unsain, Stefani, et al., 2018; Leterrier, 2021; Costa & Sousa, 2021). However, these models are not supported by direct, quantitative evidence, never mind in nerve tissue, and we speculate that a better understanding of the arrangement of spectrin tetramers along the perimeter of the axon would be instructive for understanding their function within the MPS, and the function of the MPS itself.

In this work, we investigated the organizational principles of spectrin tetramers within individual segments of the MPS in tissue. We focused our analysis on thin cryostat transverse sections of the mouse sciatic nerve, as its large and heterogeneous axonal thickness provides an ideal system to rigorously test perimetric organization *in situ*. Previous studies have shown that in myelinated fibers βII-spectrin is present in the internodes, the paranodes and the juxtaparanodes, but excluded from the nodes of Ranvier (D’Este et al., 2017). Using two-color three-dimensional dSTORM on axonal cross-sections surrounded by a thick compact myelin layer (i.e mainly internodes), we show that βII-spectrin in the MPS organizes into clusters of different sizes around the axonal perimeter. We demonstrate that βII-spectrin clusters neither follow a fully regular nor a random distribution, and that their number scales with axonal diameter, resulting in clusters consistently occupying approximately 20% of the perimeter across a wide range of axonal calibers. This work provides new insights into the axonal structure in tissue, challenging existing models of the MPS developed upon qualitative descriptions of neurons grown in culture.

## Results

### In the sciatic nerve, βII-spectrin localizes within myelinated axons and in Schwann cell somas, but is excluded from the compact myelin compartment

At low magnification, βII-spectrin immunofluorescence in nerve cross-sections highlights the characteristic concentric organization of myelinated fibers within nerves (Fig. 1A). At higher magnification, the outer ring-like structure in each fiber with strong βII-spectrin labeling corresponds to the Schwann cell soma. In contrast, βII-spectrin is absent from the compact myelin compartment of the Schwann cell, as evidenced by its lack of colocalization with the compact-myelin marker Myelin Basic Protein (MBP; Fig. 1B). In turn, the central structure of the myelin compartment surrounding the axon shows expression of βII-spectrin and the intra-axonal region displays a high concentration of βIII-tubulin (Fig. 1C). Hence, multichannel confocal microscopy reveals that βII-spectrin is mainly located in the Schwann cell soma and is also accumulated at the boundary between the compact myelin and the axoplasm. This concentric topographic feature has already been described in cross sections of nerves for both αII- and βII-spectrin (Cifuentes-Diaz et al., 2011; Einheber et al., 2013) and facilitates the identification of βII-spectrin signals arising from individual axons in single-color images. The highly irregular contour of axons observed is a widely established feature of nerves (Bremer et al., 2010; Ikeda & Oka, 2012; Yamamoto et al., 2013), and differs drastically from the quasi-circular perimeter of axons grown in culture.

**Figure 1.**
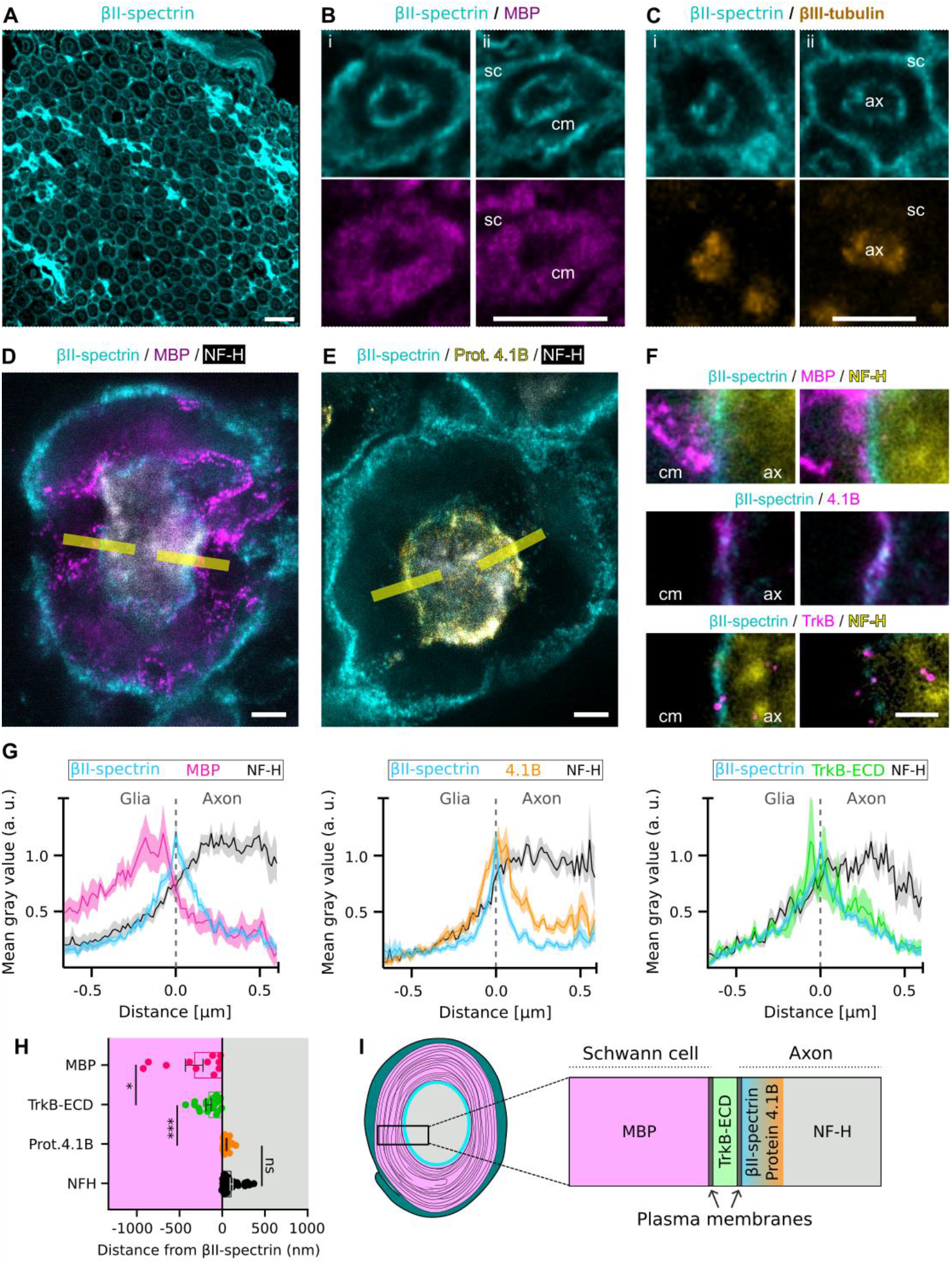
Expression and localization of βII-spectrin in cross sections of the sciatic nerve. **(A)** Low magnification confocal image of a cross-section of an adult mouse sciatic nerve immunostained for βII-spectrin (cyan). The characteristic concentric organization of myelinated axons is observed: an outer ring corresponding to the soma of the ensheating Schwann cell, an unlabeled region corresponding to compact myelin, and an inner ring surrounding the axon. Scale bar: 20 μm. **(B-C)** Magnification of individual myelinated axons double immunostained for βII-spectrin (cyan) and MBP (magenta, B) or βIII-tubulin (orange, C). βII-spectrin is located at both sides of the MBP signal, and outside of the βIII-tubulin signal. *sc*: Schwann cell soma; *cm*: compact myelin compartment; *ax*: axon. Scale bars: 5 μm. **(D-E)** STED microscopy images showing the distribution of βII-spectrin (cyan) combined with MBP (D), or protein 4.1B (E), and NF-H (gray, confocal channel). Yellow lines exemplify where intensity profiles were collected. Scale bars: 1μm. **(F)** Magnified regions of STED images exemplifying where the intensity profiles for βII-spectrin, MBP, protein 4.1B, TrkB-ECD and NF-H were obtained. *cm*: compact myelin compartment; *ax*: axon. Scale bar: 0.5 μm. **(G)** Normalized mean intensity profiles aligned to the βII-spectrin peak (arbitrary zero). Left: MBP and NF-H are distributed on opposite sides of βII-spectrin. Center: Protein 4.1B presents a profile almost identical to that of βII-spectrin. Right: The TrkB-ECD peak is slightly more external relative to that of βII-spectrin. Solid lines represent the mean, and shaded areas indicate the ± SEM. **(H)** Graph showing the linear distances between each protein to βII-spectrin. Negative values are arbitrarily located toward the exterior of the axon, and positive values toward its interior. Mean ± SEM. Pair-wise comparisons reflect one-way ANOVA followed by Tukey’s post-hoc tests (***: p<0.0001, *: p<0.05, ns: non significant). All groups were significantly different from zero (i.e. different from βII-spectrin, one-sample t test, p<0.05) **(I)** Representative diagram summarizing the inferred localization of βII-spectrin with respect to the other proteins evaluated.

The ensheating glia outer membrane and the axonal membrane are in close contact and confocal microscopy does not allow to determine whether the βII-spectrin signal observed in the axonal periphery originates from inside the axon membrane, inside the myelin sheath, or from both compartments. Previous STED analyses in longitudinal aspect of teased nerves suggested that βII-spectrin is absent in the Schwann cell side of this contact (D’Este et al., 2017), and we took a super-resolution approach in our nerve cross-sections to determine the relative position of βII-spectrin in this Schwann cell-axon contact. We performed STED microscopy combined with a quantitative analysis of intensity profiles to compare the relative spatial distribution of βII-spectrin relative to proteins with known localizations. We used MBP as a marker of compact myelin; the neuronal protein 4.1B, a marker of the cortical skeleton just below the axonal plasma, which is known to associate with actin and spectrin (Cifuentes-Diaz et al., 2011); the extracellular domain of tropomyosin kinase receptor B (TrkB-ECD) as a marker of the extracellular region of the axonal plasma membrane, expressed in a subset of sensory and motor neurons in the sciatic nerve (Foster et al., 1994; Bhattacharyya et al., 1997; Johnson et al., 1999); and neurofilament heavy (NF-H) as a marker of the axoplasm (Fig. 1D-F). The latter was always acquired in confocal mode. Intensity profiles of these proteins were analysed from lines drawn perpendicular to the glia-axon contact with accumulation of the respective markers (Fig. 1D-F, yellow lines).

To analyse the relative distance among these proteins, the intensity coordinate of each profile was first normalized to its own maximum value, and the spatial coordinate was moved so that the peak intensity of βII-spectrin corresponded to zero (Fig. 1G). We found that the MBP signal is positioned on one side of the βII-spectrin peak and the NF-H signal on the other, indicating that βII-spectrin is located between both markers, i.e., close to the membrane region (Fig. 1G, left panel). In turn, the signal from protein 4.1B reproduces the intensity profile of βII-spectrin almost identically, strongly supporting the picture that βII-spectrin localizes in the axonal cortical skeleton (Fig. 1G, central panel). The peak of TrkB-ECD is slightly towards the glial cell with respect to βII-spectrin, suggesting that the latter belongs to the axonal compartment (Fig. 1G, right panel).

For each of the profiles, we also measured the relative distance between the intensity peak of βII-spectrin and the location to the other proteins. We found that βII-spectrin is located 320 ± 100 nm from MBP (11 profiles), 155 ± 30 nm from TrkB (17 profiles), 55 ± 10 nm from protein 4.1B (23 profiles), and 115 ± 15 nm from NF-H (41 profiles) (Fig. 1H). Together, these results indicate that βII-spectrin is located within the axon, specifically on the inner face of the axonal plasma membrane (Fig. 1I). This tissue preparation allowed us to perform the next set of experiments to unravel organizational principles of βII-spectrin in individual segments of the MPS.

### 3D-dSTORM imaging of βII-spectrin reveals a non-uniform distribution along the axonal perimeter consistent with the MPS

To investigate the nanoscale organization of βII-spectrin in individual MPS segments of sciatic nerve axons, we performed two-color three-dimensional (3D) dSTORM imaging of βII-spectrin and βIII-tubulin. Transversal sections of the sciatic nerve from 4 mice were stained and imaged in three independent sessions, collecting a total of 18 full-frame regions of interest (ffROI). Axons to be further analysed within these ffROI were selected according to the following sample quality criteria. We first discarded ffROIs which could not be appropriately drift-corrected or in which the axons showed an oblique cross-section, rather than truly transversal (which was evident because βIII-tubulin signals from axons would show an elliptical shape in the same direction). Among the conserved ffROIs, axons were selected according to the following criteria: having a sharp βIII-tubulin staining and being delimited by an homogenous thick myelin sheath. After such selection, a total of 34 axons from 9 different ffROIs were included in the study. The 3D localizations confirmed the rather irregular cross sections of axons and revealed that βII-spectrin localizations are organized in groups of different sizes, mostly surrounding the βIII-tubulin localizations (Fig. 2A). To facilitate the analysis of protein localizations within segments of the MPS of sciatic nerves, we developed a custom-made open-source software that allows visualizing localizations along the *x, y* and *z* axes, and cropping and rendering the localizations of selected regions of interest. A detailed description of the software is provided in the *Materials and Methods* section.

**Figure 2.**
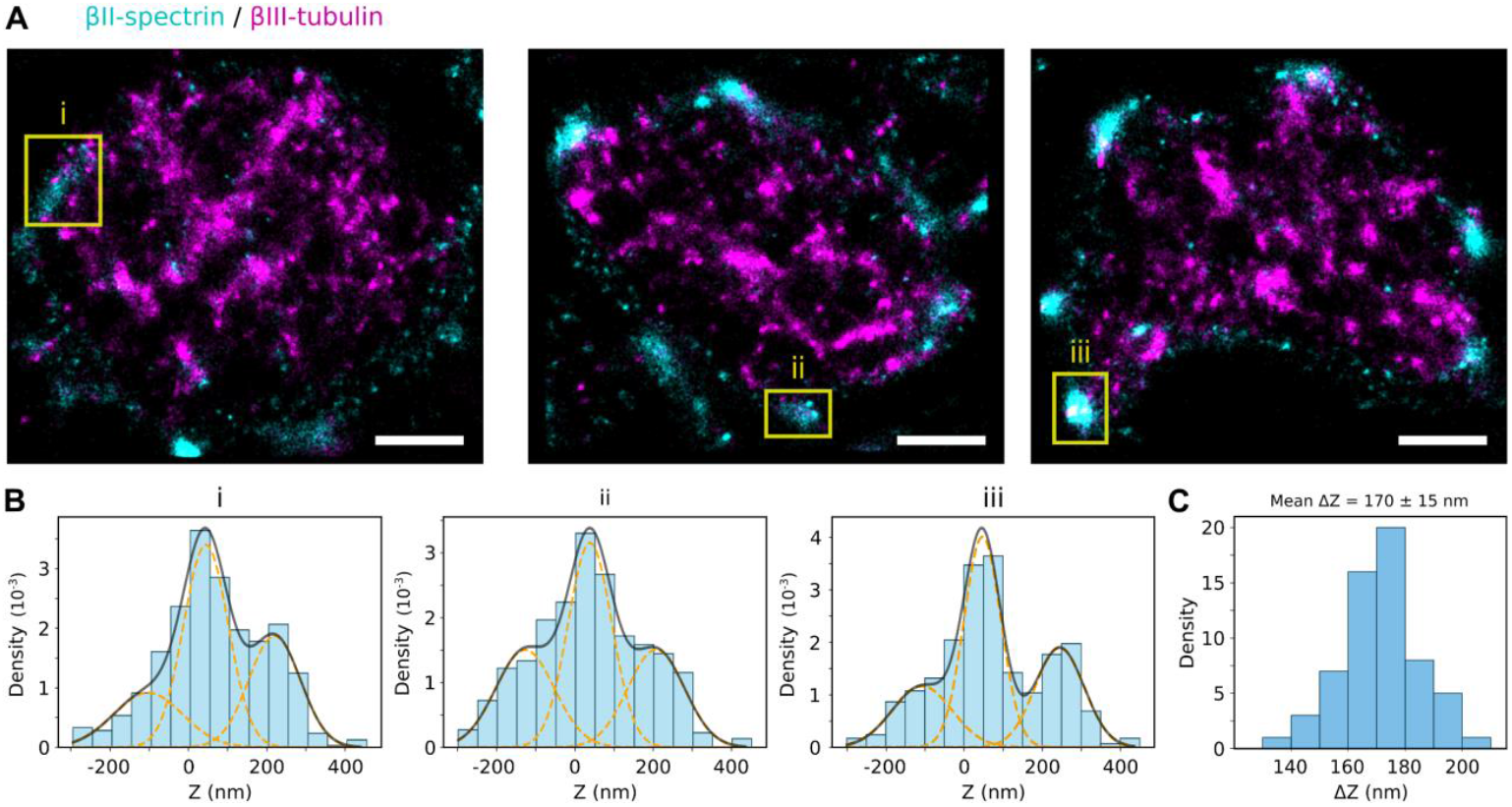
3D-dSTORM imaging of βII-spectrin reveals a non-uniform distribution along the axonal perimeter consistent with the MPS. **A)** dSTORM image of individual axons stained for βII-spectrin (cyan) and βIII-tubulin (magenta) with the z axis (∼800 nm) collapsed to show only two dimensions. Scale bars: 1 μm. **(B)** Selected regions from the reconstructed axons, indicated by orange squares in (A), used for quantitative analysis of the axial (z) distribution of localizations. Peaks in the axial localization profiles were identified by fitting a Gaussian mixture model (GMM; orange dashed curves). The inter-peak distance was calculated as the separation between the estimated means of consecutive Gaussian components. The corresponding kernel density estimation (KDE) is shown in black. **(C)** Distribution of inter-peak distances measured along the *z* axis, pooling all values obtained from the analyzed axonal cross-sections. The mean inter-peak distance measured is 170 ± 15 nm.

We first investigated whether axonal βII-spectrin exhibits the characteristic axial periodicity expected for the MPS. To this end, we analyzed the z-coordinate distributions of βII-spectrin localizations within individual regions of interest (ROIs) along axons, constrained to the axial range accessible by our 3D dSTORM measurements (∼800 nm). For each ROI, the z-distribution was modeled using a Gaussian mixture model (GMM), with the optimal number of components (2–3) selected based on the Bayesian information criterion. Only dominant Gaussian components, defined by a minimum mixture weight threshold (>5% of the z-values), were retained and sorted by their axial position. The mean positions (*μ*) of these Gaussians were interpreted as the most probable axial locations of βII-spectrin. When multiple components were detected within a ROI, the axial distances between consecutive Gaussian means (ΔZ) were computed. Three representative examples of z-distributions with the corresponding GMM fits and identified dominant components are shown in Figure 2B, corresponding to the regions highlighted in Figure 2A. Figure 2C shows the distribution of ΔZ obtained from selected ROIs from all axons. These results show that axonal βII-spectrin in peripheral nerves exhibits a periodicity of 170 ± 15 nm along the axonal axis—a value consistent with MPS measurements in other systems—strongly supporting that the βII-spectrin localizations are part of the MPS.

### βII-spectrin organizes in clusters of different sizes whose abundance scales linearly with axon perimeter

Next, we investigated whether βII-spectrin follows an organizational pattern within each segment of the MPS. For each axon analyzed, we selected βII-spectrin localizations most likely belonging to the same MPS segment by pulling localizations from a 180 nm axial range centered around the main peak of the axial localization distribution. These localizations were then analyzed with the DBSCAN algorithm (Ester et al., 1996) to identify clusters. The minimum number of localizations per cluster was set to 10 in order to match the mean number of switching cycles expected for the fluorophores used (Dempsey et al., 2011). Considering that the lateral localization precision of our dSTORM measurements was ∼20 nm, the DBSCAN searching radius ε was set to 25 nm. Figure 3A shows examples of the βII-spectrin clusters detected in three axonal sections.

**Figure 3.**
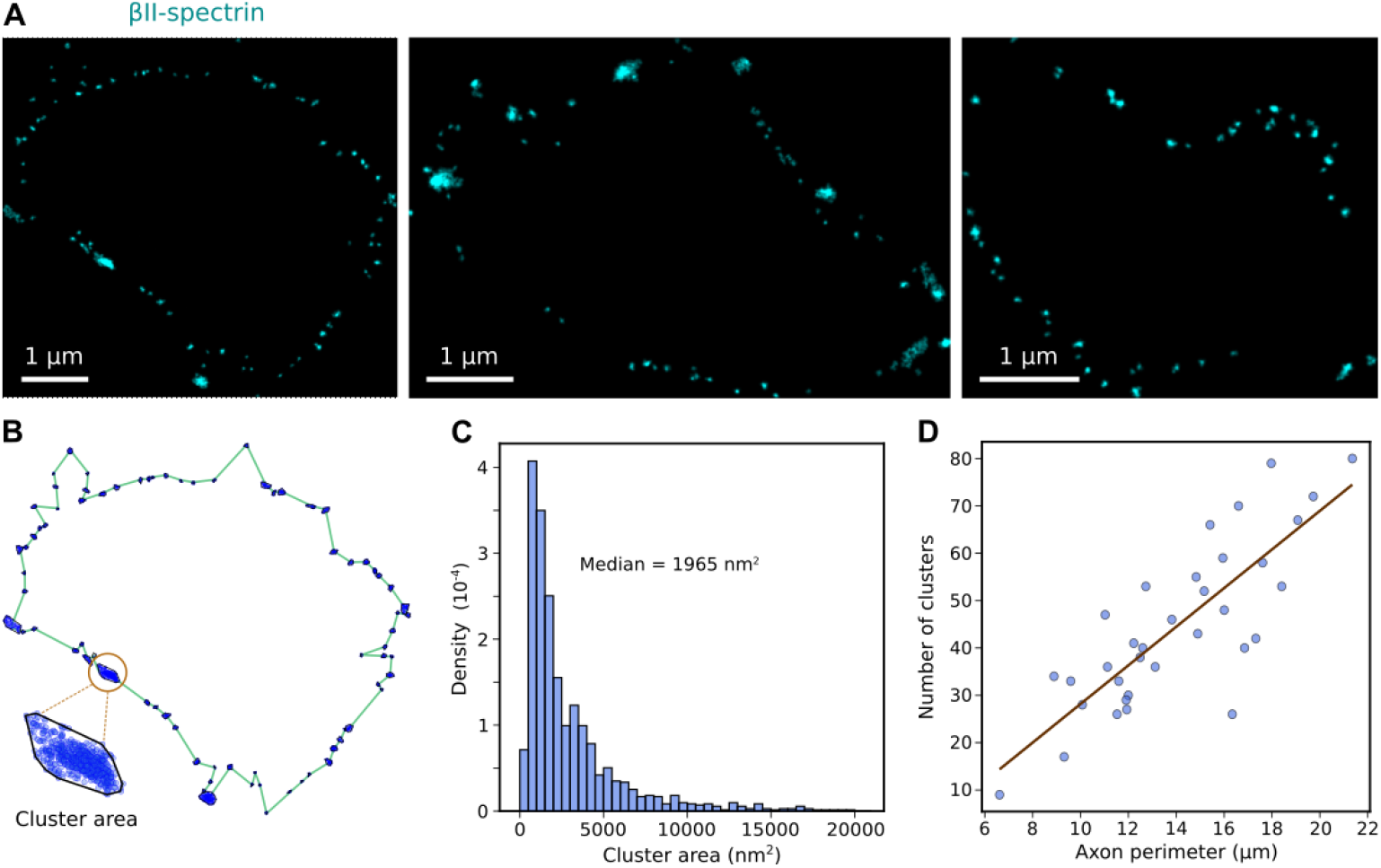
βII-Spectrin localizations form clusters of varying sizes along the axonal cortex. **A)** Example images obtained by rendering the single-molecule localizations of the βII-spectrin clusters. **(B)** Estimation of axon perimeter and cluster area. The areas shown correspond to the clusters rendered in the first example image in (A). **(C)** Distribution of βII-spectrin clusters area. Median: 1965 nm^2^ **(D)** Scatter plot of number of clusters per axon vs axon perimeter. The linear regression (y = 4.08x −12.62), shown in brown, with a Pearson correlation coefficient of 0.81, indicates a significant positive correlation between the two variables.

The axonal perimeter was reconstructed by connecting the centers of mass of the βII-spectrin clusters. Figure 3B shows an example of the reconstructed perimeter of the axon depicted in the first panel of Figure 3A. In agreement with the observation made with confocal microscopy, axons of the sciatic nerve do not maintain a circular or elliptical cross-section, unlike cultured neurons. In contrast, they present an irregularly shaped contour. The area of each cluster was estimated as the area of the smallest convex polygon containing all localizations of the cluster, as shown in the example in Figure 3B. The resulting distribution of cluster areas from the 34 different axon cross sections exhibits a pronounced peak with a median value of approximately 1950 nm^2^ (Fig. 3C), corresponding to an effective radius of about 25 nm. This size is compatible with that expected for an individual spectrin tetramer when considering the localization precision, the size of the antibodies used for immunolabeling, and the presence of two labeling sites per tetramer. Notably, this effective radius is also comparable to the lateral localization precision achieved under our experimental conditions, indicating that it represents the smallest cluster size that can be reliably resolved in our measurements. The presence of clusters with smaller apparent areas likely reflects statistical fluctuations associated with limited numbers of localizations and reconstruction effects near the resolution limit. In addition to this main population, the distribution also includes larger clusters, which were present in all axons examined, and may represent oligomeric states of spectrin tetramers. Remarkably, the number of clusters per axon scales linearly with the axonal perimeter (Fig. 3D), indicating that the number of βII-spectrin clusters increases proportionally with axonal size.

### A non-random distribution of clusters consistently covers 20% of the axonal perimeter

To investigate whether the scaling of cluster number with axon perimeter arises from a regulated distance between βII-spectrin clusters, we analyzed the nearest-neighbor (NN) distances between the centers of mass of βII-spectrin clusters displayed along the axonal perimeter of individual MPS segments. The distribution of these distances for all axons analyzed is shown in Figure 4A, which displays a prominent peak reflecting a characteristic minimal distance between adjacent βII-spectrin clusters of ∼200 nm within a given MPS segment. When examining each MPS segment individually, the medians of the NN distances remain remarkably consistent, despite differences in axonal shape and perimeter (Fig. 4B). This suggests that the minimal local spacing between βII-spectrin clusters is tightly regulated, such that axons with larger diameters incorporate additional clusters to maintain a relatively constant density. Altogether, this analysis reveals that βII-spectrin clusters follow a locally ordered but globally adaptable spatial pattern, allowing the MPS to maintain structural integrity while adapting to the morphological diversity of axons in tissue.

**Figure 4.**
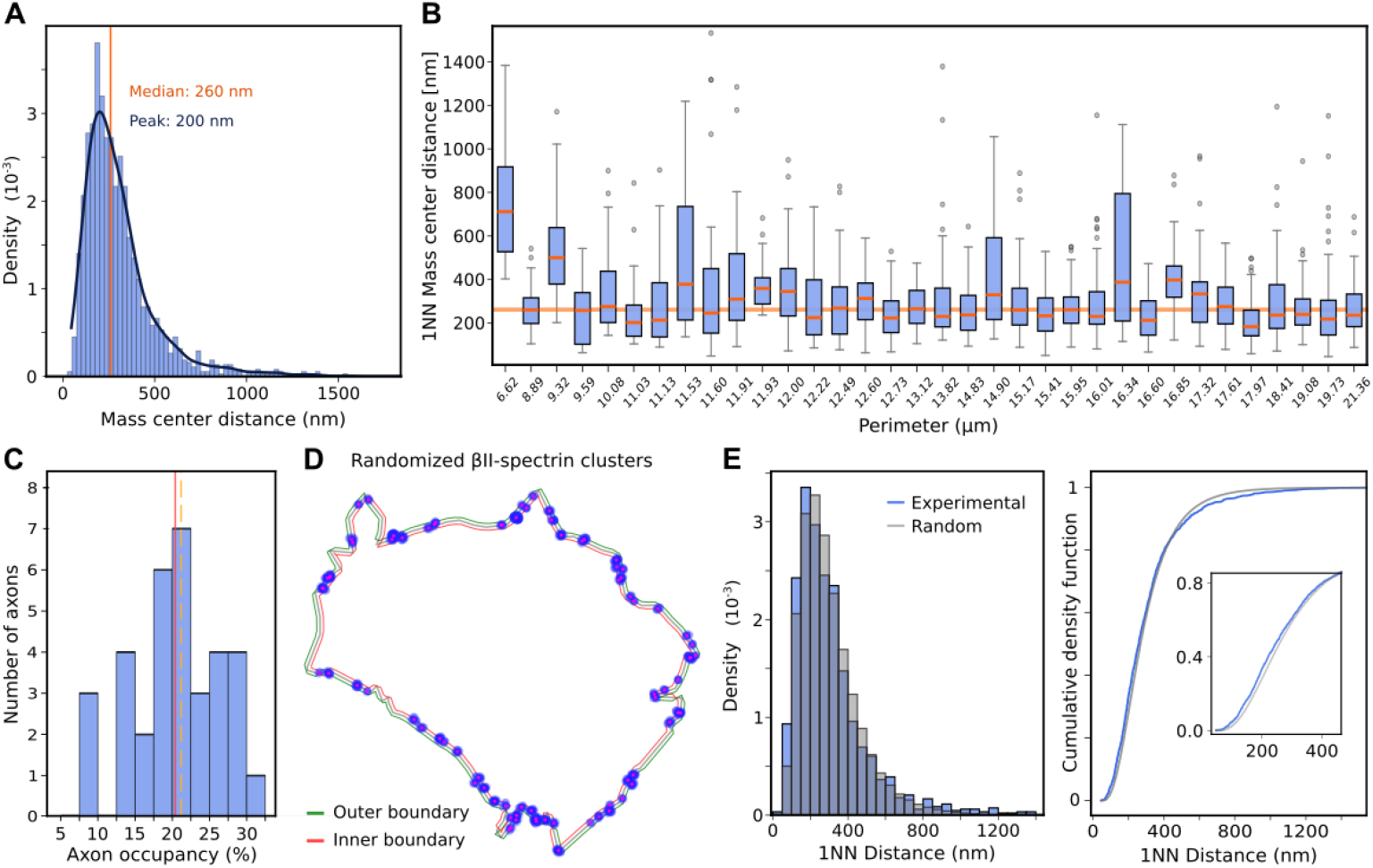
Spatial analysis of βII-spectrin clusters distribution along the axon perimeter. **A)** First nearest-neighbor (1NN) cluster distance distributions. **(B)** Boxplots for individual axons showing the relationship between axonal perimeter and the 1NN distances between βII-spectrin cluster mass centers. The median value for each axon is highlighted in orange. The horizontal light orange line in the background represents the median of all values (260 nm). **(C)** Distribution of axon occupancy values by βII-spectrin clusters across all analyzed axons. **(D)** Representative scheme of a randomized distribution of βII-spectrin clusters along an axonal perimeter. The axonal contour defined the inner (red) and outer (green) boundaries of a 100 nm-wide annular region. For each axon, cluster centers were randomly assigned within this region under a minimum inter-cluster distance constraint (based on the smallest experimental 1NN distance), with internal cluster structure (blue) preserved via rigid-body translation. **(E)** Left: histogram of 1NN distances collected from the experimental data and from randomized βII-spectrin clusters. N = 1000 randomizations were performed for each axon. Right: cumulative density functions of experimental and random distances (Inset: magnification of plot. Kolmogorov-Smirnov D = 0.054, p-value<10^-3^).

Then, we estimated βII-spectrin axon occupancy, defined as the percentage of the axonal perimeter occupied by βII-spectrin clusters. To this end, the axonal perimeter was discretized into 10,000 equally spaced points at distances < 1 nm. Each perimeter point was then evaluated to determine whether it should be considered occupied by any of the detected clusters (see *Materials and Methods* for details). Briefly, a constrained 2D Gaussian was fitted to each βII-spectrin cluster to estimate its spatial extent. Perimeter points falling within a Mahalanobis distance of 3 from the Gaussian center of the cluster were considered occupied. For each axon, we measured the cumulative length of βII-spectrin–labeled perimeter segments and expressed it as a percentage of the total perimeter. On average, βII-spectrin occupied only ∼20% of the axonal perimeter (Fig. 4C). Unlike previous suggestions from qualitative evidence in cultured neurons (Xu et al., 2013; Ganguly et al., 2015; Szalai et al., 2021), βII-spectrin distribution in MPS segments of peripheral nerves is discontinuous, with extensive stretches of the perimeter lacking βII-spectrin.

To gain insight into the particular organization of βII-spectrin within each MPS segment, we compared the experimentally observed distribution of clusters to a random distribution. To this end, we devised a randomization framework that redistributes the clusters within a 100 nm-wide region surrounding the axonal perimeter (Fig. 4D). Briefly, the axonal contour was first smoothed using B-spline interpolation, and offset curves positioned ±50 nm from this contour delineated the inner (red) and outer (green) boundaries of the region. A dense grid of candidate positions was generated, retaining only points between the boundaries. For each axon, cluster centers were randomly assigned within this region under a minimum inter-cluster distance constraint equal to the minimum first nearest-neighbor (1NN) distance measured experimentally for that axon. Then, the experimentally detected clusters were reassigned to one of the randomly generated centers via a rigid-body translation, thus preserving the internal cluster structure of each cluster. The randomization procedure was repeated 1000 times per axon. In each iteration, the 1NN distances between cluster centers were recalculated and stored (see *Materials and Methods* for details). To compare the spatial distributions of clusters in the experimental and randomized datasets, we analyzed their 1NN distance distributions (Fig. 4E). The differences are statistically significant (p-value<10^-3^), indicating that the experimentally observed spatial arrangement of clusters is unlikely to arise from a random placement along the axonal MPS perimeter. Interestingly, the *cumulative density functions* (CDFs) for the experimental and randomized distributions intersect near 0.8 (Fig. 4E, right, inset), suggesting a complex spatial organization that cannot be characterized by a simple shift toward either segregation or anti-segregation. Specifically, for approximately 80% of the cluster pairs, the experimental data exhibit a greater prevalence of shorter distances compared to the randomized model. Beyond this range, however, the experimental distribution shows relatively fewer large distances, implying spatial exclusion or regulated dispersion at larger scales.

Collectively, these observations provide evidence that the experimentally observed organization of clusters along the axonal perimeter reflects biologically regulated spatial patterning rather than a random distribution within the accessible membrane region.

## Discussion

The MPS is a highly conserved and essential cytoskeletal structure present in all neurons, particularly in axons. Yet, its molecular architecture is still not fully understood, especially in the context of native tissue. The primary objective of this work was to investigate the organizational principles of βII-spectrin within the MPS in transverse sections of the mouse sciatic nerve. Moving beyond traditional working models derived from cultured neurons, we provide new, *in situ* quantitative data on the arrangement of βII-spectrin in nerve tissue.

We first confirmed that βII-spectrin is an axonal component, present within the axon of the myelinated fibers, but specifically excluded from the compact myelin sheath. A comparative study of established markers such as MBP, TrkB-ECD, and protein 4.1B, using super-resolution STED microscopy, allowed us to localize βII-spectrin to the axonal cortical cytoskeleton—consistent with its proposed role as a structural scaffold within the MPS. Because we selected axonal cross-sections surrounded by a thick compact myelin layer, our analyses were likely performed mainly on the internodal regions of myelinated axons. An axon-specific localization of βII-spectrin in the juxtaparanodes has also been reported using longitudinally teased nerve fibers imaged with STED microscopy (D’Este et al., 2017). Additional support for this conclusion comes from the analysis of conditional βII-spectrin knockout mice. Susuki and colleagues showed that axon-specific deletion of the βII-spectrin gene eliminates the βII-spectrin signal surrounding the axon at the juxtaparanodes, demonstrating that all βII-spectrin present beneath the compact myelin sheath originates from the axon (Susuki et al., 2018). Thus, our findings also extend previous reports on βII-spectrin periodicity in intact nervous tissues (D’Este et al., 2016; He et al., 2016; D’Este et al., 2017) using a novel approach which combines nerve cross-sections and 3D super-resolution, and confirms that the axonal MPS is preserved in peripheral nerve tissue. As previously stated, βII-spectrin is absent at nodes of Ranvier—where it is replaced by βIV-spectrin (D’Este et al., 2017; Susuki et al., 2018). Future interesting studies could be conducted in order to examine whether βIV-spectrin at the nodes of Ranvier displays organizational patterns similar to those we describe here for βII-spectrin in the internode.

Utilizing 3D-dSTORM microscopy on transverse nerve sections, we determined that βII-spectrin is organized in discrete clusters that occupy only approximately 20% of the axonal perimeter. The number of clusters scales linearly with axonal perimeter, indicating that the occupancy level remains constant, irrespective of axonal dimensions. Crucially, this pattern cannot be ascribed to incomplete labeling, as the identified clusters are robustly labeled and well delineated. If βII-spectrin were uniformly distributed along the axonal circumference but at diminished density, we would anticipate a more diffuse and continuous signal rather than the distinct punctate clusters observed. Moreover, the three-dimensional nature of our measurements enabled us to verify that these peripheral clusters correspond to MPS segments, exhibiting the characteristic ∼185 nm spacing along the axonal axis. Our measured period of 170 ± 15 nm is lower than the typically reported ∼185 nm in cells in culture, yet it is congruent with previous assessments of periodicity in the sciatic nerve conducted by STED (D’Este et al., 2016), which indicated that the median periodicity in sciatic nerve tissue and retinal bipolar cells in culture was consistently ∼175 nm, whereas other cells exhibited medians of ∼190 nm. It would be intriguing to ascertain which component or organizational principle of the MPS in these disparate cells and contexts account for the variations in period observed.

A quantitative analysis of the cluster areas revealed a unimodal distribution with a predominant peak at 1,965 nm^2^. Considering the molecular dimensions of spectrin and the size added by antibody labeling, this area is consistent with that expected for a single labeled spectrin tetramer, indicating that these predominant clusters represent individual tetramers. In addition to this main population, the distribution also contains a smaller fraction of larger clusters with areas compatible with oligomeric assemblies of spectrin tetramers. The ability of spectrins to form oligomers beyond the usual dimer-to-tetramer configuration has been recently reported in developing axons of cultured hippocampal neurons (Boyer et al., 2026). Complementary techniques would be needed to determine if these larger clusters represent true oligomers (i.e. interacting tetramers) or simply non-interacting tetramers that group together non-specifically.

Mapping the center positions of βII-spectrin clusters along the axonal perimeter revealed a robust spatial organization. Within individual MPS segments, the clusters exhibit a locally ordered pattern, maintaining a consistent nearest-neighbor spacing of ∼200 nm regardless of the total perimeter length. Previous reports based on electron micrographs of purified spectrin tetramers and actin filaments have shown that the tip of the spectrin tetramer can bind with strong affinity to the actin filament in any location along the filament (Cohen et al., 1980) and not only at its ends, as commonly depicted in working models of the MPS. A comparison with a randomized distribution shows that the arrangement of βII-spectrin in the nerves deviates strongly from randomness, supporting the idea that spectrin tetramers are not associated at arbitrary positions along the actin rings and suggesting that other components of the actin ring may condition the exact location of spectrin tetramer binding. BetaII-spectrin in the MPS operates within a complex molecular environment in the axonal shaft rather than in isolation. It is known to associate with multiple components of the cortical cytoskeleton, including adducin, ankyrin-B, and protein 4.1B, both in cultured neurons and in intact nervous tissue (Baines, 2010; Bennett & Lorenzo, 2013; Xu et al., 2013; Zhou et al., 2022). Although the precise spatial organization of these interactions within the MPS remains poorly defined, the positioning of spectrin tetramers could likely be constrained by recurrent structural or molecular features of these interactions. Alternatively or in combination, constraints could arise from spectrin-anchoring complexes at the plasma membrane. Spectrin tetramers can interact firmly with the plasma membrane, both directly through multiple domains mainly located in β-spectrins (Grzybek et al., 2006), and indirectly via binding to ankyrin. Hence, protein or lipid complexes in the plasma membrane could also condition the spatial distribution of spectrin tetramers, thus potentially producing a higher-than-expected inter-cluster spacing. Although resolving the molecular determinants underlying this stereotyped inter-cluster spacing lies beyond the scope of the present study, our quantitative in situ measurements provide a framework to test whether defined spectrin-associated complexes impose geometric constraints on MPS organization in native axons.

Taken together, these findings reveal a consistent organizational principle of βII-spectrin and the MPS in sciatic nerve axons. We show that βII-spectrin occupies only a small fraction—approximately 20%— of the axonal perimeter. This limited coverage, which is consistently maintained in all axons studied regardless of axon size, may reflect a local dynamic assembly-disassembly equilibrium of βII-spectrin in segments of the MPS in native tissue. A dynamic assembly and disassembly of complete MPS segments has recently been described in axons which grow in culture (Heller et al., 2025), and highlights the possibility that in the larger myelinated axons studied here, the equilibrium state is reflected in a low but constant coverage in fixed samples.

Moreover, our study provides the first characterization of the spatial arrangement of βII-spectrin within individual MPS rings, demonstrating a non-random, locally ordered distribution with a mean nearest-neighbor spacing of ∼200 nm. This insight refines our understanding of MPS organization and challenges overly simplified structural models. By defining the precise *in situ* arrangement of βII-spectrin in healthy axons, our work also establishes a valuable reference framework for future investigations into pathological alterations of the MPS. This is particularly relevant given the involvement of spectrin and other MPS components in neurodegenerative and severe neurological disorders (Liu & Rasband, 2019; Qi et al., 2021; Stevens & Rasband, 2021; Lorenzo et al., 2023).

In conclusion, our results demonstrate that the organizational principles of the axonal MPS in native tissue are more complex, and possibly more dynamic, than previously appreciated. The clustering behavior of βII-spectrin and its scaling with axon size point to an adaptable cytoskeletal architecture. These findings underscore the importance of quantitative, *in situ* measurements in tissue samples to guide accurate working hypotheses about the structure and function of the MPS *in vivo*.

## Materials and Methods

### Animals

C57BL6 mice were born in the vivarium of INIMEC-CONICET-UNC (Córdoba, Argentina). C57BL6 mice lines were originally provided by Charles River Laboratories International Inc (Wilmington, USA). All procedures and experiments involving animals were approved by the Animal Care and Ethics Committee (CICUAL) of INIMEC-CONICET-UNC (Resolution numbers 014/2017 B, 015/2017 B, 006/2017 A and 012/2017 A) and were in compliance with approved protocols of the National Institute of Health Guide for the Care and Use of Laboratory Animals (SENASA, Argentina).

### Reagents

The following primary antibodies were used in this study: anti-βII-spectrin (mouse, 1:400, BD Biosciences cat. #612563); anti-MBP (myelin basic protein, rabbit, 1:500, generously provided by Dr Ana Lis Moyano -CIMETSA-IUCBC, Argentina-(Landry et al., 1996); anti-NF-H (neurofilament heavy, chicken, 1:1000, Abcam cat. #ab4680); anti-βIII-tubulin (rabbit, 1:1000, BioLegend cat. #PRB-435P);TrkB-ECD (extracellular domain, rabbit, 1:500, Millipore Cat. #07225), anti-protein 4.1B (rabbit, 1:400, generously provided by Dr Laurence Goutebroze -Sorbonne Université, France (Cifuentes-Diaz et al., 2011)). Secondary antibodies used for confocal and STED microscopies were: anti-mouse-IgG-STAR ORANGE (goat, 1:250, Abberior cat. #STORAGE-1001); anti-rabbit-IgG-STAR ORANGE (goat, 1:250, Abberior cat. #STORAGE-1002); anti-rabbit-IgG-STAR RED (goat, 1:250, Abberior cat.#STRED-1002) and anti-chicken-IgY-AlexaFluor488 (confocal only, donkey, 1:500, Jackson Immunoresearch cat. #703-545-155). Secondary antibodies used for dSTORM microscopy were: anti-mouse-IgG-Alexa Fluor 647 (goat, 1:500, Invitrogen, Cat # A-21235); anti-mouse-IgG-Alexa Fluor 568 (goat, 1:500, Invitrogen, Cat # A-11011).

### Sciatic nerve sections

#### Transcardiac perfusion fixation

Four P60 male C57BL6 mice underwent transcardiac perfusion to fix the animal’s tissues. Briefly, animals were anesthetized with an intraperitoneal injection of 6% chloral hydrate (0.5 ml/100 g). Once fully anesthetized, the heart was exposed, the cannula was inserted into the left ventricle, and an incision was made in the right atrium. At this point, the peristaltic pump was started and washing began (0.4% glucose, 0.8% sucrose, 0.8% NaCl, 0.04% heparin –for a total of 2000 IU). Once no blood was observed, the solution was changed to the fixation solution (0.005% Na_2_SO_3_, 0.38% Na_2_B_4_O_7_, 4% PFA [Sigma-Aldrich, cat. no. 441244], 1% H_3_BO_3_). The animal was fixed for at least 15 to 20 minutes.

#### Dissection and post-fixation

For sciatic nerve dissection, the animals were placed in ventral recumbency. The body of each mouse was sprayed with 70% ethanol. An incision was made in the upper part of the leg using a small No. 15 scalpel, taking care not to cut too deeply and limiting the incision to separating the muscle fibers to expose the nerve. In all cases, only the segment of the sciatic nerve prior to the bifurcation to the tibial nerve was collected. The collected tissues were placed in vials containing 4% PFA and left overnight at 4°C.

#### Sectioning

To perform cryoprotection, tissue was submerged for 24 hours in each of the following sucrose solutions: 10%, 20% and 30% in PBS. Once cryoprotection was complete, the tissues were embedded in Cryoplast (Biopack, Argentina) using plastic molds to control tissue position relative to the cryostat blade. The resulting blocks were stored at –80°C until further processing. Tissue sections 5 µm thick were made in a Leica CM1850 cryostat. The sections were mounted on previously gelatinized slides or coverslips (1% gelatin, Pura Química, Argentina), according to the microscopy technique to be used. Finally, the preparations were left to dry at room temperature overnight and stored at –20°C until use for immunofluorescence staining.

### Immunofluorescence

The samples were washed twice in PBS for 5 minutes. They were then blocked with a solution of 0.1% Triton X-100 (Cicarelli, Argentina) and 5% horse serum (HS) in PBS for 1 hour at room temperature. The primary antibodies were incubated in 0.1% Triton X-100 and 2.5% HS overnight at 4°C. Three washes were performed with PBS for 5 minutes. The secondary antibodies were incubated for 2 hours at room temperature in the same solution. When staining for myelin basic protein (MBP), a blocking and antibody incubation solution with 2% Triton X-100 was used. Samples were mounted in Mowiol (2.4% Mowiol 4–88 (poly(vinyl alcohol), Sigma) when confocal or STED imaging was performed, or in dSTORM buffer when imaging was performed using this technique.

### Confocal microscopy

Microscopy images were acquired at CEMINCO (Centro de Micro y Nanoscopía de Córdoba, Argentina, https://ceminco.conicet.unc.edu.ar/) using a Zeiss LSM800 inverted microscope (AxioObserver platform) equipped with a Plan-Apochromat 63x/1.40 NA oil immersion objective. The images were captured using internal photomultiplier tube (PMT) detectors and acquired using ZEN software (version 14.0.27.201). Alexa Fluor 488, 568, and ATTO 647 dyes were sequentially excited using 488, 561, and 633 nm lasers, respectively. Emission was detected with spectral PMTs in ranges of 493–598 nm, 568– 682 nm, and 638–759 nm. Z-axis slices were acquired with a step size of 0.3 µm and a final voxel size of 0.13 × 0.13 × 0.3 µm. Image averaging (2×) was applied during acquisition. Imaging conditions were configured to avoid saturated pixels and allow observation of linear changes in intensities. All imaging parameters were kept constant across all conditions. Images were subsequently processed and analyzed (as indicated below) using Fiji (ImageJ), with brightness and contrast adjusted uniformly for visualization only; no additional processing was applied.

### Stimulated Emission Depletion Nanoscopy

Stimulated emission depletion (STED) nanoscopy images were acquired at CEMINCO (Centro de Micro y Nanoscopía de Córdoba, Argentina, https://ceminco.conicet.unc.edu.ar/) using a STEDYCON STED super-resolution microscope (Abberior Instruments, Germany), installed on an Olympus IX81 inverted microscope using a UPlanXApo 100X oil immersion objective, NA:1.45. For two-color STED, the samples were excited with a 580 nm laser and a 635 nm laser, and depleted with a 775 nm laser. The images were obtained in a single plane with the pinhole open to ∼1 Airy unit (more information on STED acquisitions with this instrument can be found on the supplier’s website).

#### Glial-axon contact analysis

using Fiji (ImageJ) software, intensity profiles were obtained for all channels along the glial-axon contact, using the βII-spectrin signal as a reference. Intensity profiles were saved and subsequently processed in Excel. Briefly, the background was subtracted and the profiles were smoothed using the running average method. The smoothed signals from βII-spectrin, 4.1B and TrkB were mostly Gaussian-shaped, and a normal Gaussian fit was used to find the position of their peaks. The peak of βII-spectrin was set to zero. The signal from MBP and NF-H in most cases had a geometric growth from βII-spectrin, and then reached a plateau (instead of a peak). For these cases, to calculate their distance to βII-spectrin, we considered the distance to the first value that was 20% lower than the maximum. Displayed STED images were smoothed and contrasted using ImageJ for the purposes of better visualization of the final figure, but quantifications were performed on raw images.

### 3D Stochastic Optical Reconstruction Microscopy

#### dSTORM sample preparation

Sciatic nerve sections were mounted on previously gelatinized 22 mm coverslips and placed in a holder. Imaging was performed in dSTORM imaging buffer (50mM Tris, 10mM NaCl, 10% w/v glucose, pH=8) supplemented with 1 μg/mL glucose oxidase (Sigma-Aldrich), 0.5 ug/mL catalase (Sigma-Aldrich) and 10 mM MEA (mercaptoethylamine) as oxygen scavenging system.

#### Imaging

The dSTORM microscope was custom-built around a commercial inverted microscope stand Olympus IX-73 operating in wide-field epifluorescence mode. A 642 nm 1.5 W laser (MPB Communications 2RU-VFL-P-1500–642) and a 532 nm 1.5 W laser (Laser Quantum Ventus 532), both circularly polarized, were used for fluorescence excitation. A 405 nm 50 mW diode laser (RGB Photonics Lambda Mini) was used for reactivating fluorescent molecules. The lasers were focused to the back focal plane of the oil immersion objective Olympus PlanApo 60x NA 1.42. Collected fluorescence light was decoupled from the laser excitation by a quad-edge multiband mirror (Semrock Di03-R405/488/532/635-t1). Further blocking of the illumination lasers was performed with a quad-notch filter NF (Semrock NF03-405/488/532/635E-25). Dual-color simultaneous imaging was performed through emission wavelength discrimination. A dichroic lowpass filter (Chroma ZT647rdc) and two band-pass filters (Semrock 582/75 BrightLine HC and Chroma ET700/75m) were used to separate the fluorescence emission from Alexa Fluor 568 and Alexa Fluor 647, respectively. Three-dimensional super-resolution image acquisition was performed using astigmatic detection. A cylindrical lens (CL, Thorlabs LJ1516RM-A, f = 1000 mm) was positioned between the tube lens and the camera, introducing a controlled astigmatism that encodes the axial position into the shape of the point spread function (PSF) without significantly compromising lateral resolution.

#### Calibrations

Prior to dSTORM measurements, a 3D PSF calibration dataset was acquired using *Dark-Red* beads (Life Technologies FluoSpheres™ carboxylate-modified microspheres 0.04 μm) at a concentration of 10^−6^ M. The 642 nm laser was operated at 1–3 W cm^−2^. The beads were brought into focus without active focus stabilization, as an axial scan was performed across the sample. A Z-stack was recorded with a step size of 25 nm over a total axial range of 2000 nm (±1000 nm), generating the calibration curve required for precise axial localization. Differences in magnification, shear and image rotation between the two channels were corrected prior to acquisition by imaging multicolor beads (Life Technologies Tetraspeck 0.1 μm), performing an affine transformation correction for the overlay of channels (Hartley & Zisserman, 2003/2003). The emission light was expanded with a 2x telescope so that the pixel size of the EMCCD camera (Andor iXon3 897 DU-897D-CS0-#BV) was 133 nm in the sample plane. The camera and lasers were controlled with Tormenta, a custom software developed in the laboratory (Barabas et al., 2016).

#### Image acquisition

Prior to dSTORM imaging, conventional fluorescence images of the regions of interest (ROIs) were acquired using an excitation power density of 1–3 W cm^−2^. Channel overlay calibration was performed before each acquisition round to ensure accurate alignment between both detection channels. Super-resolution data collection was then initiated by increasing the excitation laser power density to 10–20 kW cm^−2^, promoting stochastic on–off switching of the fluorophores. Throughout the whole acquisition, the 405 nm laser power was increased whenever the density of single-molecule blinking events decreased below 1 molecule per μm^2^. Each acquisition consisted of approximately 16,000 frames with an exposure time of 50 ms per frame.

### Image post-processing

Post-processing was carried out using the *Picasso* software package (Schnitzbauer et al., 2017), where single-molecule localization was performed using the maximum likelihood estimation (MLE) Integrated Gaussian method (Smith et al., 2010). Localizations were filtered based on the global localization precision (12nm) estimated by the NeNA algorithm (Endesfelder et al., 2014). *Picasso* was also used for rendering the super-resolution images. Additionally, redundant cross-correlation (RCC) drift correction was applied to all datasets (in each one, we first corrected the drift via RCC in one of the emission channels to generate a drift-correction file, which was then applied to correct the drift in the second channel to ensure that both channels had the exact same correction).

### MPS Explorer

Axon selection was performed using the interactive environment *MPS Explorer* (https://github.com/luhalac/MPS-explorer), a tool specifically developed for the exploration and analysis of 3D-dSTORM data. This platform enabled simultaneous handling of multiple channels, 2D/3D visualization of signals, and precise delineation of each axon within the acquired volume. The analysis pipeline comprised: (1) loading the data in.hdf5 or.csv formats; (2) multichannel visualization and overlay of βII-spectrin and βIII-tubulin signals; (3) selection of subregions of interest using graphical tools; (4) axial filtering based on defined Z-coordinate ranges; (5) automatic detection of molecular clusters using the DBSCAN algorithm (Ester et al., 1996); (6) computation of inter-cluster distances and generation of histograms to identify potential periodic patterns; (7) removal of low-quality clusters; and (8) final export of the curated results. This selection and curation stage enabled the generation of a representative dataset of axons with high-quality signal, in which the three-dimensional organization of the MPS could be analyzed with both axial and perimetral resolution.

### Statistical analysis

#### STED intensity profiles

Data are presented as mean ± SEM and most graphs show also the individual values of the replicates from which the mean and SEM were calculated, as indicated. Calculations were made using the statistical software GraphPad Prism 10. Normality and homoscedasticity were tested for each group in order to apply the following parametric tests. When comparisons were made among three or more groups, one-way ANOVA was used; followed by Tukey post-hoc multiple comparisons, to find where the differences occurred. Significance is indicated with asterisks when p ≤ 0.05 and detailed in each case. To assess whether the distances of the proteins were different from the location of βII-spectrin, i.e. different from zero, we performed a one-sample t test.

#### DBSCAN

Density-Based Spatial Clustering of Applications with Noise (Ester et al., 1996) was applied to the lists of STORM localizations obtained. This clustering method was chosen due to the unknown number and shape of the clusters and the presence of isolated localizations in the data (noise). MinPoints parameter was set to a value of 10, following the mean number of switching cycles expected for Alexa Fluor 568 and Alexa Fluor 647 (Dempsey et al., 2011).

#### Axon coverage estimation

The percentage of coverage was estimated as the proportion of the axonal perimeter occupied by βII-spectrin clusters. To this end, the axonal perimeter was discretized into 10,000 equally spaced points at distances < 1 nm. Each perimeter point was then evaluated to determine whether it should be considered occupied by any of the detected clusters. For each cluster, we computed the mean and covariance matrix of the (x, y) coordinates and fitted a Gaussian distribution whose maximum standard deviation was limited to one third of the largest distance from the cluster center to any of its points. This constraint ensured that the fitted Gaussian accurately reflected the true spatial extent of the cluster without overestimating its spread. Using the inverse covariance matrix, we then calculated the Mahalanobis distance from each perimeter point to the cluster center. Points lying within a Mahalanobis distance of 3σ_max_ were considered to be “occupied” by that cluster, where σ_max_ corresponds to the largest standard deviation of the fitted ellipses.

#### Cluster randomization

To assess whether the spatial distribution of clusters along the axonal membrane deviates from randomness, we developed a spatial randomization framework that redistributes clusters within an annular region surrounding the axonal perimeter. First, the perimeter of each experimental axon was smoothed using B-spline interpolation to generate a continuous contour. Offset curves were then computed at fixed distances of ± 50 nm from this contour, thereby defining the inner and outer boundaries of the annular region. This resulted in a 100 nm-wide crown-like domain encircling the axon, which approximates the spatially accessible membrane surface area. The selected width accounts for the estimated membrane thickness (∼5–7 nm) and the combined localization uncertainty arising from the point spread function and antibody size (∼20–30 nm). Within this annular region, each experimental cluster was translated as a rigid body to a new random position, without overlap and while respecting a minimum inter-cluster distance that was not smaller than the smallest first nearest-neighbor (1NN) distance observed in the experimental data. The randomization procedure was repeated 1000 times per axon. In each iteration, the 1NN distances between cluster centers were recalculated and stored, generating a null distribution of spatial relationships for each axon.

The study design was not pre-registered. No randomization was performed to allocate subjects in the study. No sample size calculation was performed. Outliers were identified, and excluded from analysis, using the ROUT method (Q=1%), implemented through GraphPad Prism 10.

## Ethics approval

All procedures and experiments involving animals were approved by the Animal Care and Ethics Committee (CICUAL) of INIMEC-CONICET-UNC (Resolution numbers 014/2017 B, 015/2017 B, 006/2017 A and 012/2017 A) and were in compliance with approved protocols of the National Institute of Health Guide for the Care and Use of Laboratory Animals (SENASA, Argentina).

## Consent for publication

Not applicable.

## Availability for data and materials

All data generated or analyzed during this study are included in this published article. The raw data and analyses can be shared upon request to the corresponding authors

## Competing interests

The authors declare that they have no competing interests.

## Funding

ANPCyT-PICT-2021-GRF-II-00048.

## Author contributions

N.G.G., A.M.S., F.D.S. and N.U. contributed to the conceptualization of the study by formulating the overarching research goals and aims. The methodology, including the development and design of experimental approaches, was established by N.G.G., G.E., L.F.L., E.A.G., A.M.S. F.D.S. and N.U. N.G.G., G.E., L.F.L., A.M.S., F.D.S. and N.U. validated the replication and reproducibility of results, experiments, or other research outputs. Software development, programming, implementation of computer code or algorithms, including testing code components was performed by L.F.L., G.E. and A.M.S. The formal analyses and investigation phase, which included performing experiments and data collection, was undertaken by N.G.G., G.E., L.F.L., A.M.S. and N.U. Resources, including the provision of materials, instrumentation, and analysis tools, were contributed by L.F.L., L.G., M.B., E.A.G., A.M.S., F.D.S. and N.U. Data curation was performed by G.E., L.F.L. and A.M.S. The original draft of the manuscript was written by N.G.G., G.E., F.D.S. and N.U., while all authors contributed to review and editing. Visualization of data and preparation of figures were carried out by N.G.G., G.E., F.D.S. and N.U. Supervision including oversight, leadership responsibilities and mentorship were carried out by M.B., E.A.G., A.M.S., F.D.S. and N.U. Project administration, including coordination and execution of the research activities, was led by F.D.S. and N.U. Funding to support the study was secured by M.B., F.D.S. and N.U.

## Acknowledgements

N.U. acknowledges research funding from ANPCyT, Argentina (PICT-2021-GRF-II-00048). The authors greatly acknowledge the technical and imaging assistance of Dr. Gonzalo Quassollo, Dra. Cecilia Sampedro, Dr. Carlos Mas and Dr. Pilar Crespo from Centro de Micro y Nanoscopía de Córdoba – CEMINCO – CONICET – Universidad Nacional de Córdoba, Córdoba, Argentina. N.G.G. and G.E. had PhD fellowships from CONICET during the execution of this work.

## Notes

### Competing Interest Statement

The authors have declared no competing interest.

